# From Fill to Freeze: Redesigning homeRNA to Enable Robust Sampling and Scalable Downstream Processing Workflows for Remote Blood Transcriptomics

**DOI:** 10.64898/2026.11.04.764386

**Authors:** Madeleine Eakman, Ingrid H. Robertson, Xiaofu Wei, Filip Stefanovic, TJ Caira, David J. Guckenberger, Karen N. Adams, Erwin Berthier, Lauren G. Brown, Sanitta Thongpang, Ashleigh B. Theberge

## Abstract

In-person research can introduce barriers to participation, including proximity to a study site and scheduling constraints. Our lab developed the homeRNA platform to address these barriers, enabling remote blood collection and RNA stabilization for transcriptomic research. To increase the applicability of homeRNA for large-scale studies, a redesigned version was developed with modifications to 1) improve usability for both participants and downstream processing laboratories, 2) allow for high-throughput manufacturing, and 3) extend the shelf life of the stabilizer tube. In this work, we systematically tested this design against anticipated stresses that can occur throughout a remote study (e.g., high temperature exposure during shipment) to identify and characterize its limitations. We found that the homeRNA device can be prepared and stored at room temperature at least three months in advance of sampling with continued shelf life analysis underway. We also found that the homeRNA platform produces RNA of sufficient quality for downstream transcriptomic analysis under standard shipping durations (i.e., 2 days in transit) and ambient-to-high temperature conditions (i.e., 25-40 °C) and that RNA quality is unaffected by variable blood volumes collected by participants (mean RIN=7.8 in blood volumes 0.25-1 mL). Lastly, we also found that samples can be stored in typical cold storage temperatures and even up to standard room temperature for at least 4 weeks (mean RIN=8.1 after 4 weeks of storage at -80 to 20 °C). This work validates that the new homeRNA platform is robust to many of the expected stresses that samples can experience during remote studies.

## Introduction

The human body can have complex and dynamic biological responses to pathogens, toxins, ongoing disease progression, or other stimuli over time, but the specific mechanisms for these responses are still being investigated. Blood is useful for measuring a wide range of analytes (e.g., RNA, DNA, proteins) in response to such exposures or disease states^1–4^, but collecting blood samples for such studies has historically relied on in-person venous draws, which can create barriers to participating, such as participant proximity to a clinic, scheduling conflicts, and access to transportation^5^. These barriers are exacerbated in studies requiring frequent sampling timepoints. At-home studies can reduce these barriers by eliminating the need for a participant to travel to a study site and allowing for more frequent sampling on a more flexible schedule. Dried blood spot (DBS) samples have been successfully used for transcriptomics profiling^6,7^, but common stresses in remote studies (e.g., exposures to heat and humidity) can accelerate RNA degradation in these samples during shipping^8^.

Our lab has developed a growing suite of tools to better enable remote transcriptomics research. Specifically, we have developed homeRNA^9–11^, which typically comprise an upper arm capillary blood collection device and a custom-designed stabilizer tube, allowing for blood collection and subsequent RNA stabilization at home. Thus far, homeRNA has been shown to preserve RNA samples in real-world^12^ and in-lab^13^ controlled high-temperature exposures and has been used in several longitudinal transcriptomic studies^14–17^. Our lab recently developed homeRNAmax^10^, which is compatible with the commercially available BD Microtainer widely used across various upper-arm blood collection devices (e.g., Tasso+, RedDrop, TAP) and allows for the collection and stabilization of up to 1 mL capillary blood.

Importantly, our previous longitudinal transcriptomics studies involved enrollment on the order of 40-100 participants with ≤20 samples each. To design larger scale remote studies with hundreds to thousands of participants and samples, it is important to have a platform that can scale accordingly. Our prior homeRNA stabilizer tubes required a time-consuming gluing step that limited kit production to a maximum of a few hundred stabilizer tubes in a week. The homeRNA tube therefore had to be redesigned to allow manufacturing of thousands to tens of thousands of kits to empower larger studies. For example, the stabilizer tube design was simplified to parts that twist together, completely removing the need for gluing when assembling homeRNA kits. To ensure that this new design can effectively enable large-scale remote studies in a reproducible way, we needed to stress test the new platform across all the different scenarios that are possible in remote sampling.

A typical study using homeRNA platforms is executed in four phases (Fig. 1): (1) kit preparation, where a technician fills a homeRNA tube with RNAlater stabilizing solution; (2) sample preparation, where a participant collects capillary blood from the upper arm using a lancet-based blood collection device and stabilizes the RNA using homeRNA; (3) sample shipping, where the sample is shipped back to the lab for analysis; and (4) sample storage, where the stabilized sample is held until extraction and analysis. In an “ideal” scenario, homeRNA tubes would be filled with RNAlater the same day as kits are sent to participants, the participant would collect the maximum possible blood volume (1 mL), the sample would be shipped in an ambient temperature and humidity with minimal additional shaking, and the sample RNA would be either immediately extracted or the samples would be frozen at -80 °C until extraction and analysis. However, these ideal scenarios are highly impractical to real-world study designs and conditions; samples can often be subjected to delayed shipping times, high temperature exposures, or variable storage temperatures depending on available laboratory resources. To date, there has been work that characterizes RNA degradation in blood samples with changing storage temperatures and storage times^18–22^, and work that investigates how previous homeRNA platforms perform in high temperatures^12,13^; the present manuscript builds upon these prior efforts by systematically introducing stresses to the homeRNA platform and blood samples stabilized using the platform. These stresses include aging of homeRNA tubes through long-term storage prior to stabilization; reduced volumes of blood stabilized using the device; controlled vibrations, high temperatures, and low- and high-humidity incubation after stabilization; and long-term storage of blood samples at different temperatures. By testing these parameters, this work will inform guidelines and recommendations for treatment of homeRNA kits and samples in future studies.

**Figure 1.**
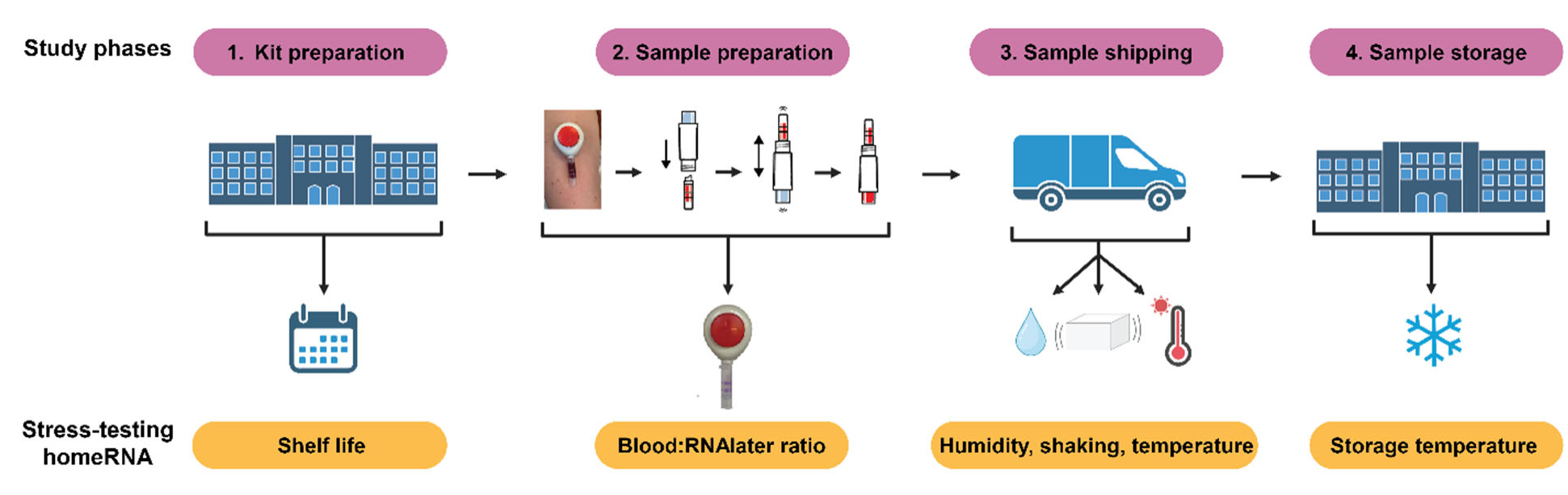
Schematic of the phases of homeRNA studies. The phases are indicated in the top (shaded pink), and the corresponding experiments performed in this work to test limitations at these phases are indicated on the bottom (shaded orange). Created using BioRender.com.

## Experimental

### Fabricating homeRNA tubes

The homeRNA adapters were injection molded out of polypropylene and sealed with thermal foil (Seabright LLC). The day of blood sample stabilization (or tube storage for the experiment assessing homeRNA tube aging, see below), sample tubes (Axygen, Corning) were filled with 3.6 mL RNAlater (Thermofisher Scientific), and labels were applied assigning conditions (e.g., low, ambient, or high humidity). After filling the sample tubes, the homeRNA adapter with thermal foil seal is then twisted onto the sample tube to completely seal the device. Thermal foil seals are removed before blood stabilization.

### Participant recruitment for venous blood draws

Venous blood draws were performed under a protocol approved by the University of Washington Institutional Review Board study number STUDY00014133. To be eligible for the study, participants had to be (1) at least 18 years old or older, (2) not pregnant, and (3) self-reported as healthy. Participants were recruited via word of mouth in Seattle, Washington, United States. Written informed consent was obtained from all participants prior to blood draws.

### Assessing the impact of homeRNA tube aging on RNA integrity (shelf life)

Thirty-six homeRNA tubes were filled with 3.6 mL RNAlater (Thermofisher Scientific). Eighteen tubes were placed in a 25 °C incubator, and eighteen tubes were placed in a 40 °C incubator for aging. After one month and three months of storage, nine tubes from each incubator were removed and used to stabilize 1.0 mL whole blood from a single donor using dipotassium ethylenediaminetetraacetic acid (K_2_EDTA) coated BD Microtainer (BD) tubes; nine additional homeRNA tubes were made the same day as blood stabilization of aged tubes to serve as a control comparison. After blood stabilization, three tubes from each storage condition (i.e., aged at 25 °C, aged at 40 °C, and freshly prepared) were incubated at 25 °C for 48 h, three tubes were placed in a 40 °C incubator for 48 h, and three tubes were immediately frozen at -80 °C as a control. After incubation, RNAlater-stabilized blood tubes were immediately stored at -80 °C until extraction and analysis. Samples were collected across two timepoints from two distinct donors (i.e., one donor for homeRNA tubes after one month of storage and another donor for homeRNA tubes after three months of storage), with all replicate samples (n=3 replicates per incubation conditions) derived from the same respective donor. One replicate from each condition was extracted in separate extraction batches with nine samples each (3 extractions, each with nine samples for a total of 27 samples at one month and 27 samples at 3 months). One month samples were extracted in the same batches, and three month samples were extracted in the same batches separate from the one month samples.

### Assessing whether excess RNAlater (i.e., blood:RNAlater ratio) impacts RNA integrity

K_2_EDTA coated BD Microtainer tubes were filled with 0.25 mL, 0.5 mL, or 1.0 mL of blood collected from a single venous draw, with 4 replicates made for each volume, to simulate blood collection variability observed with upper-arm capillary blood collection events. Filled BD Microtainer tubes were then connected to homeRNA tubes filled with 3.6 mL RNAlater and shaken to stabilize until all fluid was the same color. Two tubes of each volume were then incubated for 48 hours at 25 °C, and two tubes of each volume were immediately frozen at –80 °C as a control. After incubation, samples were immediately stored at -80 °C until RNA extraction and analysis. All samples for this experiment were extracted in the same extraction batch (twelve samples total).

### Assessing the impact of shaking on RNA integrity

K_2_EDTA coated BD Microtainer tubes were filled with 1.0 mL of venous blood, connected to homeRNA tubes filled with 3.6 mL RNAlater, and shaken to stabilize until all fluid was the same color. Three tubes were placed in an orbital shaker/incubator set to 50 rpm and 25 °C for 6 h, three tubes were placed in an incubator set to 25 °C for 6 h (no shaking), and three tubes were immediately placed into a -80 °C freezer as a control. After incubation, samples were immediately stored at -80 °C until RNA extraction and analysis. All blood samples were taken from a single venous draw. Note: This experiment was conducted at the same time as humidity experiment and shares an immediately frozen control; replicates were also extracted in the same batches (e.g., replicate 1 for the vibration experiment and replicate 1 for the humidity experiment were extracted at the same time along with replicate 1 of the immediately frozen control; replicates 2 and 3 were extracted in two additional separate batches totalling six samples for each extraction batch).

### Assessing the effect of humidity on RNA integrity

K_2_EDTA coated BD Microtainer tubes were filled with 1.0 mL of venous blood. Filled BD Microtainer tubes were then connected to homeRNA tubes filled with 3.6 mL RNAlater and shaken to stabilize until all fluid was the same color. Three tubes were placed in a Ziploc bag filled with 454 g Drierite desiccant (Thermofisher Scientific) in a 25 °C incubator to simulate low humidity (approximately 10% humidity), three tubes were placed in a 25 °C incubator with a water bath to simulate high humidity (approximately 90% humidity), three tubes were placed in a 25 °C incubator at ambient humidity (approximately 40% humidity), and three tubes were immediately stored in a -80 °C freezer. Samples were incubated for 48 h and immediately stored in a -80 °C freezer until RNA extraction and analysis. All blood samples were taken from a single venous draw. Note: This experiment was conducted at the same time as vibration experiment and shares an immediately frozen control; replicates were also extracted in the same batches (e.g., replicate 1 for the vibration experiment and replicate 1 for the humidity experiment were extracted at the same time along with replicate 1 of the immediately frozen control; replicates 2 and 3 were extracted in two additional separate batches, totalling six samples for each extraction batch).

### Assessing the impact of high temperatures on RNA integrity

K_2_EDTA coated BD Microtainer tubes were filled with 1.0 mL of venous blood. Filled BD Microtainer tubes were then connected to homeRNA tubes filled with 3.6 mL RNAlater and shaken to stabilize until all fluid was the same color. homeRNA-stabilized blood tubes were then placed in a 25 °C, 35 °C, 40 °C, and 45 °C incubator (n=9 tubes per temperature condition). Three tubes were immediately stored in a -80 °C freezer. After 24 h, 48 h, and 5 days, three tubes were removed from each incubator and stored in a -80 °C freezer until RNA extraction and analysis. All blood samples were taken from a single venous draw. Samples were extracted in three batches of 13 samples each, with one replicate from each condition included in one batch.

### Assessing the impact of sample storage conditions on RNA integrity

K_2_EDTA coated BD Microtainer tubes were filled with 1.0 mL of venous blood, connected to homeRNA tubes filled with 3.6 mL RNAlater, and shaken to stabilize until all fluid was the same color. homeRNA-stabilized blood tubes were then either placed on a benchtop at ambient temperature (approximately 20 °C, ranging from 16 to 22 °C) or stored in a 4 °C refrigerator, -20 °C freezer, or -80 °C freezer (n=6 tubes per storage condition). Six Axygen tubes (Axygen (Corning)) without the homeRNAmax adapter and six 5 mL microcentrifuge tubes (MTC Bio) that are certified for –80 °C storage were also filled with 3.6 mL RNAlater and used to stabilize 1 mL whole blood as additional controls for comparison. Three tubes were removed from each storage condition after 1 week and 4 weeks of storage and immediately stored in a -80 °C freezer until RNA extraction and analysis. All blood samples were taken from a single venous draw. Samples were extracted in 3 batches of six samples each, with one replicate from each condition included in one batch.

### RNA extraction and analysis

RNA was extracted in a method similar to that used in Eakman, Stefanovic et. al 2026^10^: in brief, samples were extracted using the standard RiboPure RNA Purification Kit protocol (Invitrogen). The samples were split into two aliquots and extracted at the same time. Prior to the elution step, these two aliquots were combined into a single sample on the same column. Resulting RNA integrity was analyzed using the Agilent 2100 Bioanalyzer Nano Kit (Agilent Technologies, Inc.) and RNA yield was analyzed using the Qubit Flex Fluorometer (Thermofisher Scientific). RNA yield for the experiment assessing blood:RNA ratio was measured using the Cytation 5 instrument (BioTek).

## Results and Discussion

### homeRNA stabilizer tube was redesigned to broadly improve usability

The original homeRNA stabilizer tube^9^ was created to be a user-friendly tool for stabilizing RNA in blood collected with the Tasso-SST blood collection device. homeRNA was initially designed in three separate injection-molded parts: a sample vial to hold liquid RNA stabilizer, an adapter to connect the sample vial and the blood tube specific to the Tasso-SST device, and a custom-designed cap to prevent RNA stabilizer evaporation and spillage during shipping. The adapter and sample vial pieces had to be glued together, leaving a nozzle opening that allowed the tube to be filled and emptied using a plastic Pasteur pipette, while also further preventing leakage and spillage upon inversion (more detail on the mechanism for this spill-resistant nozzle feature can be found in Eakman, Stefanovic et. al 2026^10^). The stabilizer tube was designed to accommodate 500 µL blood – the maximum allowable volume of the Tasso-SST blood tube – and enough RNA stabilizer to stabilize 500 µL blood, as well as additional RNA stabilizer as a safeguard.

When the Tasso-SST was discontinued, we modified the stabilizer tube design to improve compatibility with upper arm blood collection devices^10^. Instead of interfacing with a device-specific blood tube, homeRNAmax was designed to interface with the commercially available BD Microtainer blood tube, allowing homeRNA to be usable with other capillary blood collection devices beyond Tasso (e.g., RedDrop One (RedDrop Dx), TAP MicroSelect (YourBio)). BD Microtainers can hold up to 1 mL blood; the stabilizer tube volume was also therefore increased to accommodate the additional blood and stabilizer.

Through these design considerations, our previous homeRNA platforms have enabled user-friendly at-home blood collection and RNA stabilization for use in remote transcriptomics studies. However, multiple inefficiencies in previous stabilizer tube designs were not compatible with manufacturing for larger-scale remote longitudinal studies, particularly for studies that required hundreds to thousands of samples; for example, manually gluing together three separate components comprising a single homeRNA tube was time consuming and is not suited to large-scale manufacturing requirements.

Building upon the general designs of the previous homeRNA platforms, the redesigned homeRNA platform improves manufacturability, usability, and downstream processing without modifying the sampling workflow (Fig. 2). First, to remove the time-consuming gluing step required by both previous stabilizer tubes, the updated design changes the sample tube with a commercially available transport tube (Axygen), which interfaces with a custom screw-on adapter composed of a single injection molded part. In addition to the advantages of removing the gluing step, this modification also allows laboratory technicians to fill the stabilizer tube with a specific stabilizer (in our case, RNAlater) and remove the stabilized sample with a standard pipette instead of a plastic Pasteur pipette. The Axygen tube also comes in multiple volumes, allowing homeRNA platforms to be more broadly compatible with stabilizers that might require a higher stabilizer:blood ratio. Next, the custom-designed cap of the previous homeRNA platforms was replaced with a thermal foil seal, which effectively prevents evaporation of RNAlater (Fig. S1-S2) and can allow for easier removal by participants with limited dexterity compared to previous homeRNA stabilizer tube caps. Lastly, once homeRNA-stabilized samples are received in a central laboratory, the screw-on adapter can be removed for smaller footprint cold storage, increasing the compatibility of homeRNA with large-scale storage or biobanking needs. Since homeRNA also uses a standard commercially available sample tube, this tube can also be compatible with automated liquid handling setups. Together, these modifications make homeRNA tubes easier to work with for both laboratory technicians and study participants, which is vital for successful large-scale remote studies.

**Figure 2.**
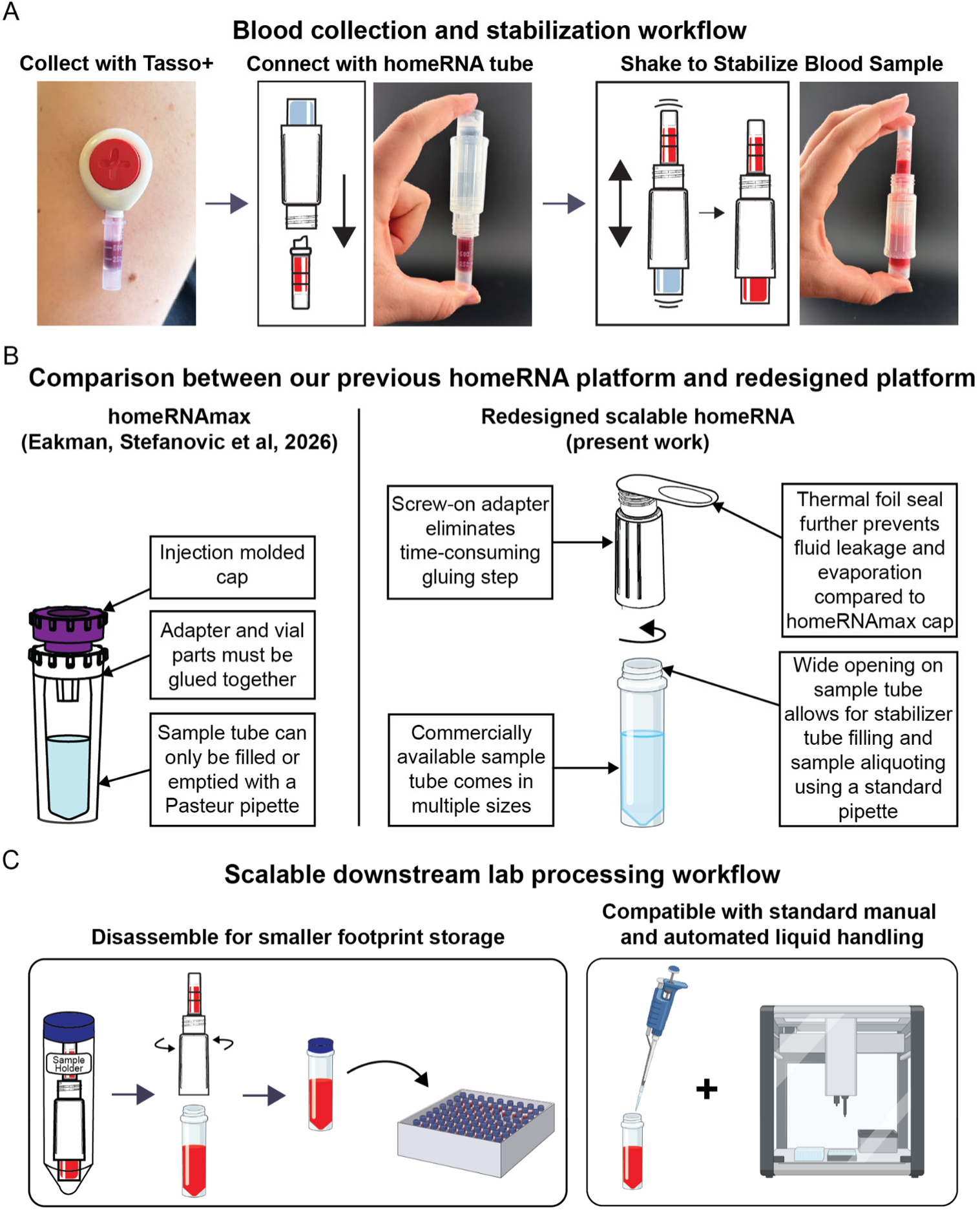
homeRNA-based sampling and stabilization workflow with stabilizer tube design modifications. A) Typical blood sampling and subsequent RNA stabilization workflow for homeRNA. Users collect up to 1 mL of capillary blood from their upper arm using the Tasso+ blood collection device into a BD Microtainer, which is then removed and connected to the homeRNA stabilizer tube containing the RNA stabilization solution. To stabilize the blood sample, users then vigorously shake the blood and stabilizer fluid together until it is the same color throughout. B) Comparison of the stabilizer tube used in previous designs of homeRNA (left; homeRNAmax) with the redesigned scalable homeRNA stabilizer tube used in this work (right). C) The redesigned homeRNA tube is compatible with downstream processing workflows, in which the homeRNA adapter can be removed to store samples within the sample tube with a smaller footprint. The sample tube is also compatible with standard laboratory pipettes or with standard automated liquid handling setups.

### homeRNA tubes can be prepared up to 3 months prior to blood stabilization with little impact to RNA integrity

To evaluate the shelf life of a filled homeRNA tube, we conducted an experiment to test RNA integrity of stabilizer tubes stored in controlled environments for extended periods of time (1 month and 3 months) (Fig. 3A). We chose to age the tubes at 25 °C to represent a typical laboratory room temperature that homeRNA tubes might be stored in, as well as 40 °C to accelerate aging. We then stabilized 1.0 mL of venous blood using these tubes, as well as tubes that were prepared the same day as blood stabilization as a control. These stabilized samples were then stress-tested through incubation at 25 °C and 40 °C for 48 hours, representing a typical return shipping time for homeRNA samples after at-home sampling.

**Figure 3.**
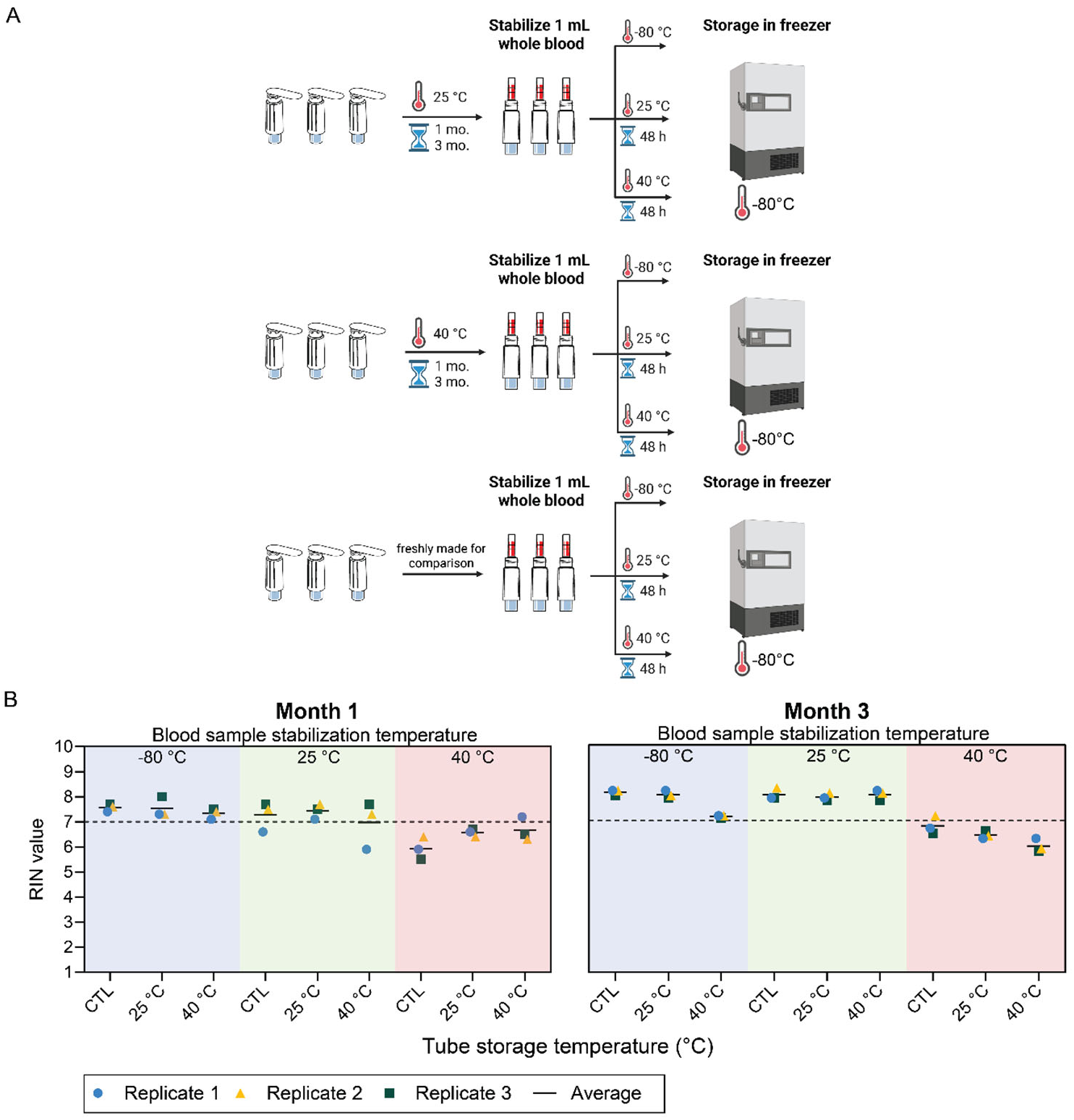
homeRNA tubes stored for three months at 25 °C and 40 °C are still able to consistently stabilize blood. A) Outline of experimental conditions. Tubes were filled with RNAlater then stored for one month and three months at either 25 °C or 40 °C, then used to stabilize 1 mL venous blood. For each storage (1 month and three months), n=3 tubes were incubated at 25 °C for 48 hours, 40 °C for 48 hours, or immediately frozen at -80 °C. For each timepoint, a set of homeRNA tubes was freshly prepared and stabilized with 1 mL of venous blood at the same time as the aged tubes to serve as a control comparison. Created with BioRender.com. B) RNA quality of each extracted blood sample. Different symbols represent replicates extracted from the same batch. Each condition was performed in triplicate from one donor at each blood stabilization timepoint. Bar represents the average of the three replicates.

We used the RNA Integrity Number (RIN), a common metric used to measure the quality of RNA samples, to measure the effectiveness of homeRNA to properly stabilize blood RNA. RIN values were measured using an Agilent Bioanalyzer 2100, which uses a proprietary algorithm to assign integrity values ranging from 1 to 10, in which a RIN of 10 represents completely intact RNA and a RIN of 1 represents completely degraded RNA^23,24^. We highlight a RIN of 7 throughout this work as a conservative cutoff suitable for the majority of downstream transcriptomic analyses, but sample RINs lower than 7 can still be usable in some applications^25–28^; for example, we used a RIN of 6 as a cutoff for RNA sequencing in a recent longitudinal study^14^. For a full list of experimental conditions, RIN, and yield data for this experiment, see Table S1.

We found that homeRNA tubes that were stored at 25 °C and 40 °C for 1 and 3 months prior to blood stabilization had RIN values similar to that of tubes that were prepared fresh the same day as blood stabilization (Fig. 3B). There is some variability between the sample RINs for a given condition (in particular, the samples that were stored at 25 °C after 1 month of storage and stabilization); batch to batch variation in extractions can lead to small variances in the RIN, but this variability is not generally meaningful. Notably, all stabilized blood samples incubated at 40 °C had RIN values less than 7. It is expected for RNA to experience some degradation after constant exposure to high temperatures, as can be seen in our prior work^12,13^. Importantly, blood samples incubated at 40 °C for 48 hours using freshly made homeRNA tubes, tubes stored for three months at 25 °C, and tubes stored for three months at 40 °C, were 6.8, 6.4, and 6.0, respectively. This shows that three months of homeRNA tube storage at elevated temperatures does not strongly affect the platform’s ability to stabilize blood, but rather the temperature that the stabilized blood samples experience prior to RNA extraction and analysis. For more information on the impact of high temperature incubation of blood stabilized using the homeRNA platform in this work, see *homeRNA samples are robust to variable humidity, excess shaking, and some exposures to high temperatures*.

### RNA integrity of homeRNA-stabilized blood samples is not affected by blood:stabilizer ratio

There are multiple reasons why it is important to understand whether excess RNAlater impacts sample quality. Firstly, the manufacturer’s recommended ratio of blood:RNAlater is 1:2.6; homeRNA tubes are filled with 3.6 mL of RNAlater for 1 mL of blood in case a participant collects more blood than instructed, or in case there is any evaporation during the shipping process. Secondly, there can be variation in how much blood participants will collect when sampling. In the Instructions for Use (SI figure), participants are instructed to either collect blood to 1 mL, or to collect blood for 5 minutes, whichever comes first. It is therefore possible that a participant collects <1 mL when they stop collecting, further increasing the ratio of RNAlater:blood in a sample. Indeed, in previous studies with both homeRNA and homeRNAmax we have seen blood collection with the Tasso devices vary from approximately 0.1-1 mL^9,10,12,14,15,17^.

In addition to the variable blood volumes that can be collected, we also filled homeRNAmax tubes with additional RNAlater as a safeguard against leakage/spillage in transit or additional drops of blood that can be collected. We therefore sought to understand any impacts that excess RNAlater and variation on collected blood volume can have on homeRNA-stabilized samples. To test different ratios of blood:RNAlater, venous blood was stabilized using homeRNA in volumes of either 0.25 mL, 0.5 mL, or 1 mL using the typical volume of RNAlater in homeRNA tubes, 3.6 mL (Fig. 4A). These samples were then incubated at 25 °C for 48 hours to represent standard shipping conditions for homeRNA. The RNA integrity remained comparable when lower volumes of blood were stabilized with the same volume of RNAlater and incubated (Fig. 4B); the mean RIN across all volumes was 7.9 after immediate storage at -80 °C and 7.8 after incubation at 25 °C for 48 hours. The homeRNA platform is therefore shown to be resilient to variable blood collection volumes, further systematically validating it for at-home use. We note that this experiment only validates homeRNA’s use with RNAlater as its stabilization solution; if another solution were to be chosen, then similar testing would need to be done to determine acceptable blood collection volumes for that solution. For more detail on the RIN and yield data for this experiment, see Table S2.

**Figure 4.**
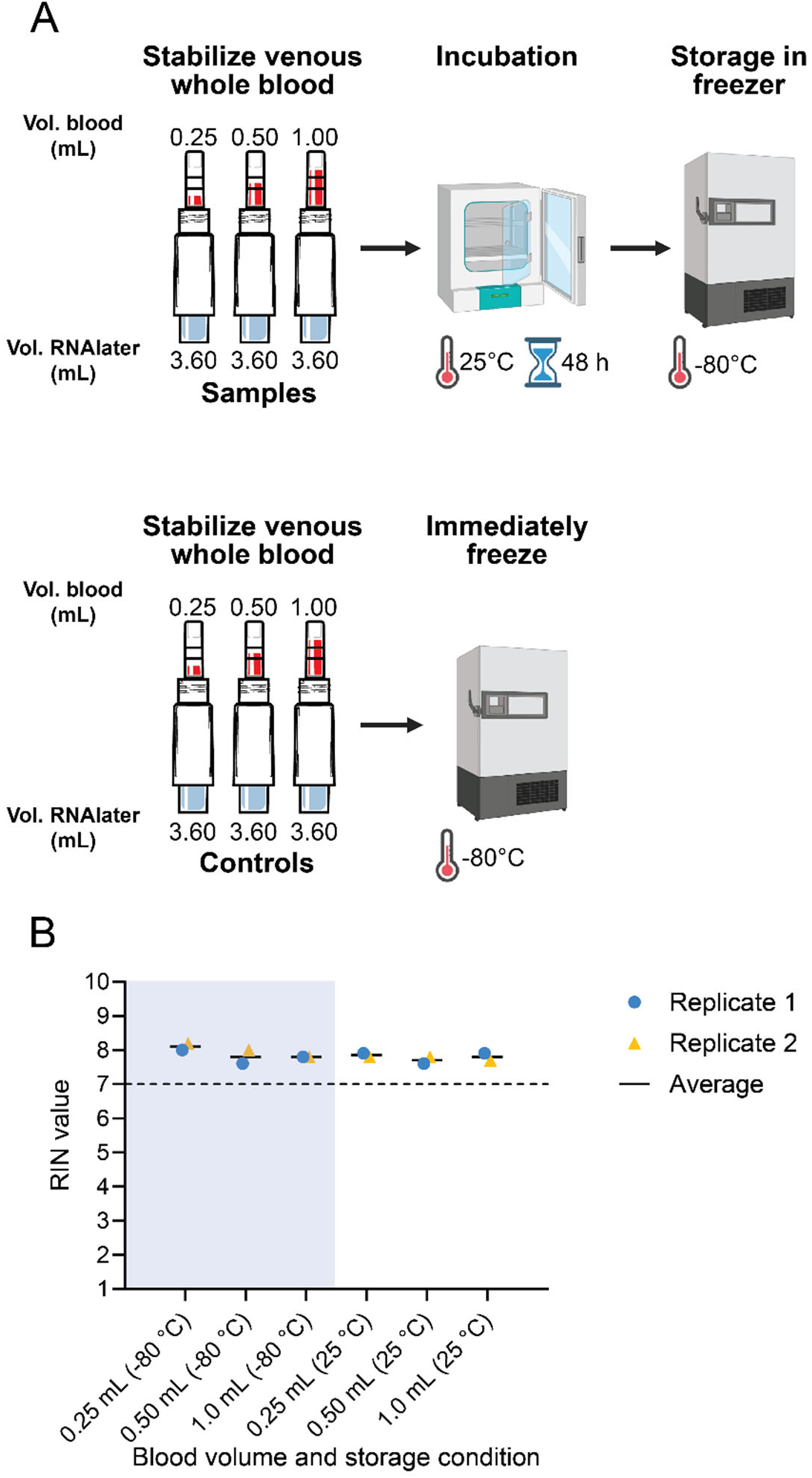
High RIN values are observed across a range of blood:RNAlater ratios. A) Schematic of experimental conditions. homeRNA tubes filled with 3.6 mL RNAlater were used to stabilize 0.25 mL, 0.5 mL, or 1.0 mL venous blood and incubated for 48 hours at 25 °C. Additional samples were also immediately frozen at -80 °C after blood stabilization as a control. Each condition was conducted in duplicate using blood from the same venous draw. Created with BioRender.com. B) RNA integrity of extracted samples. Different symbols represent different replicates. Replicates for this experiment were extracted and analyzed in the same batch. Bars represent the average RIN for each ratio and temperature across all replicates.

### homeRNA samples are robust to variable humidity, excess shaking, and some exposures to high temperatures

Samples shipped in remote studies can be subject to many weather conditions and variable treatment in transportation. We therefore tested sample stabilization when exposed to additional shaking, as homeRNA-stabilized blood samples may experience additional vibrations or disturbances in return shipment, as well as extreme humid or dry conditions and high temperatures in arid and tropical climate environments where homeRNA could be deployed.

To test whether additional vibrations and shaking impacts RNA quality in blood samples stabilized using homeRNA, samples were shaken on an orbital shaker for six hours at 50 rpm (Fig. 4A). This length of time and rate of shaking were chosen to represent more shaking than would be expected in any shipping scenario. RNA integrity was found to be comparable between shaken (mean RIN=8.8) and unshaken (mean RIN=8.8) samples after 6 hours (Fig. 5B).

**Figure 5.**
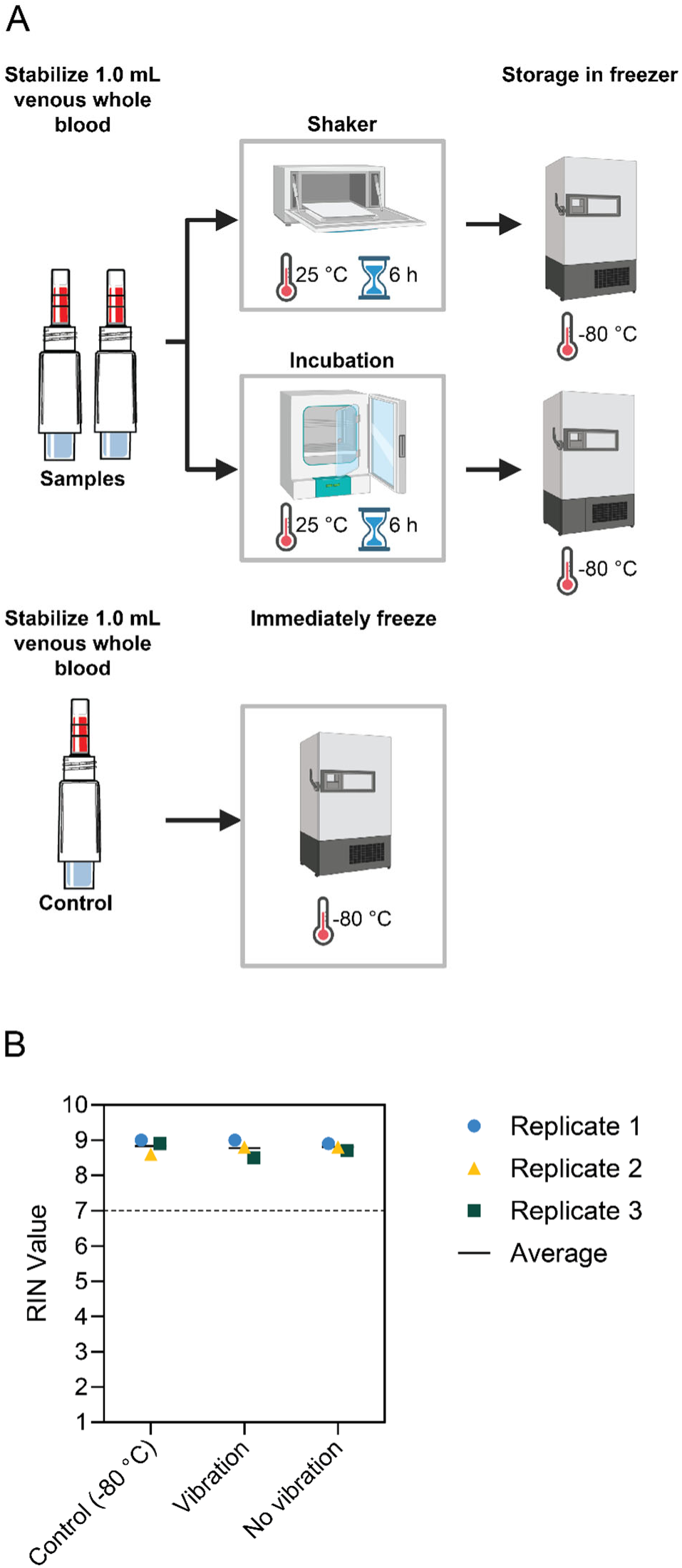
High RIN values are observed regardless of shaking. A) Schematic of experimental conditions. homeRNA tubes filled with RNAlater were either shaken for 6 hours (vibration), incubated in a 25 °C incubator for 6 hours (no vibration), or immediately frozen at -80 °C (control). Created using BioRender.com. B) RNA integrity of extracted homeRNA-stabilized blood samples. Experiment was conducted in triplicate from one donor; different symbols represent replicates and bar represents the average of the 3 replicates.

Humidity is an important factor in other remote blood sampling methods, especially in dried blood spot (DBS) sampling, where humidity has been shown to have a negative impact on resulting RNA integrity^8^. In typical homeRNA studies, homeRNA-stabilized blood samples are stored within a sealed 50 mL conical vial as a secondary containment and then shipped back within the original packaging, so we would expect humidity to have a limited impact on the sample storage and resulting RNA integrity. To test the effect of humidity on homeRNA-stabilized blood samples, we incubated samples in an extremely low and high humidity environment (10% and 90% humidity, respectively) to represent a standard shipping duration in extreme humid and dry conditions (Fig. 6A). We incubated all samples at 25 °C to represent standard temperatures in shipping. In these humidity conditions, RNA integrity was found to be comparable at ambient (33-49%) (mean RIN=8.7), extreme low (mean RIN=8.6), and extreme high humidity (mean RIN=8.6) while at 25 °C for 48 hours when compared to the control samples that were stabilized and frozen at -80 °C (mean RIN=8.83) (Fig. 6B).

**Figure 6.**
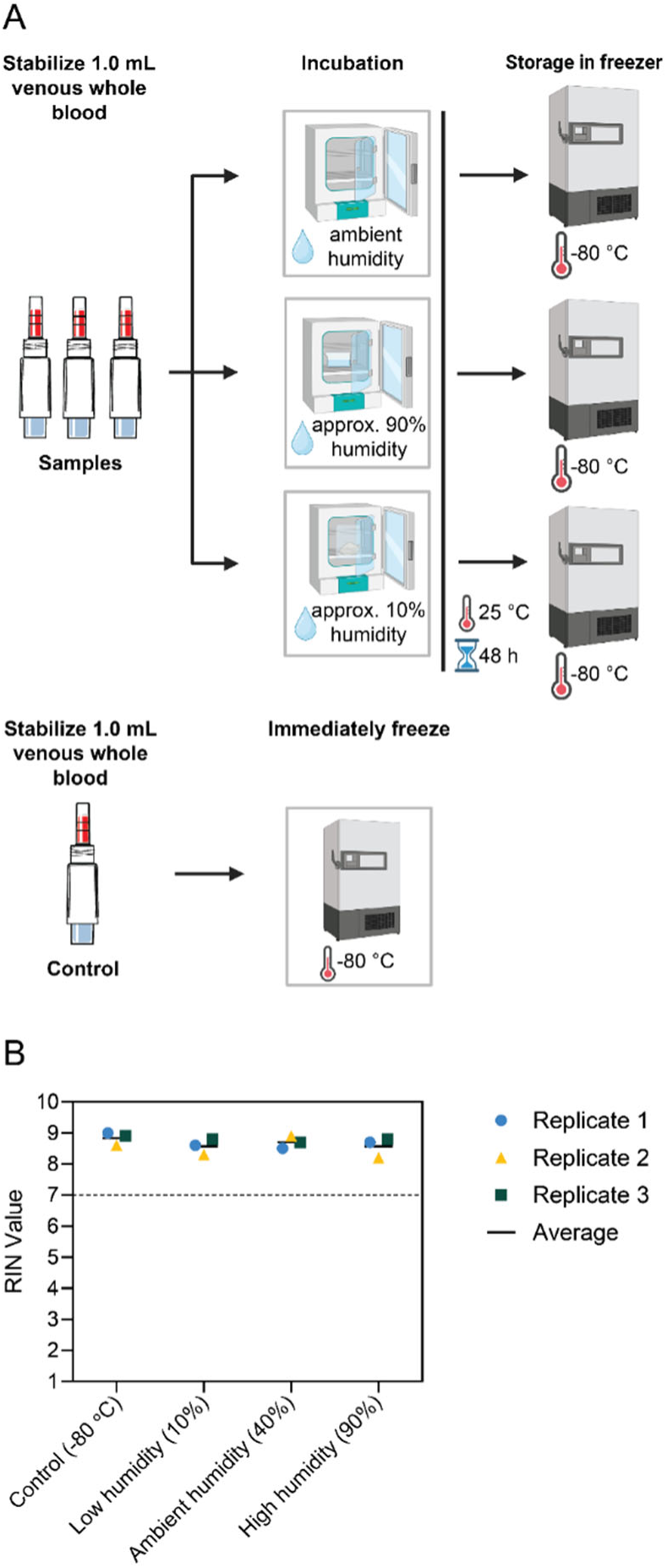
High RIN values are observed across a range of humidities. A) Schematic of experimental conditions. homeRNA-stabilized blood samples were placed in ambient, extreme low, or extreme high humidity while at 25 °C for 48 hours. Additional blood samples were also immediately frozen at -80 °C as a control. Created using BioRender.com. B) RNA integrity of extracted samples. Each condition was conducted in triplicate; different symbols represent replicates and the bar represents the average of the 3 replicates.

Finally, it is important to understand the impact of high temperature on blood samples stabilized using the homeRNA platform, as samples might be subject to high temperatures while awaiting pickup or in transit. It is worth noting that our lab has already extensively characterized our original homeRNA platform’s performance in high temperature climates and in controlled high temperature conditions^12,13^. However, the experiments in these studies were either in real-world conditions that experience variable temperature exposure at different time scales or they used samples stabilized and incubated in standard lab tubes instead of a homeRNA platform; as yet no controlled temperature-stress experiment has been conducted on homeRNA-stabilized blood samples.

For the characterization of the new design of homeRNA, we stabilized venous blood samples using homeRNA devices and incubated them at consistent temperatures representing standard and extreme temperatures that might be encountered during shipping, including 25 °C, 35 °C, 40 °C, and 45 °C. We incubated them for 24 hours and 48 hours to represent standard return shipping durations, as well as 5 days to represent a more extreme shipping duration (Fig. 7A). We found that 24 hours of temperature exposure across all conditions had little effect on RNA integrity (i.e., all had RIN values greater than 7). Additionally, even 48 hours exposure to temperatures up to 40 °C resulted in high RNA quality (RIN values greater than 7). However, after 48 hours exposure to 45 °C, RIN values dramatically dropped (mean RIN=5.4) and then continued to decrease after 5 days (mean RIN=3.0). While 25 °C exposure for 5 days was still able to properly stabilize samples (mean RIN=8.4), RIN decreased for 35-45 °C exposure after 5 days (mean RIN 6.6 and 3.0 for 35°C and 45°C, respectively); however the samples incubated at 35°C for 5 days still had an average RIN value close to 7, which could still be used for downstream transcriptomic analysis. RNA integrity decreased after incubation at 45 °C for 2 and 5 days as well as 40 °C for 5 days; these findings are expected as constant exposure to high heat has been shown to accelerate RNA degradation^8,18,20^. However, our lab’s prior work characterizing heat exposure in shipping shows that constant exposure to temperatures as high as 40-45 °C for periods as long as 2-5 days would not be expected in real-world shipping conditions, which would likely only experience exposures to >30 °C on the order of hours rather than days^13^. Additionally, our prior work shows that RNAlater-stabilized blood can be exposed to high temperatures (up to 50 °C) for hours at a time and still result in RNA of sufficient integrity for analysis (mean RIN=6.4)^13^. For summary data of experimental conditions, RIN, and yield for the humidity, shaking, and high-temperature experiments in the present manuscript, see Tables S3-S4.

**Figure 7.**
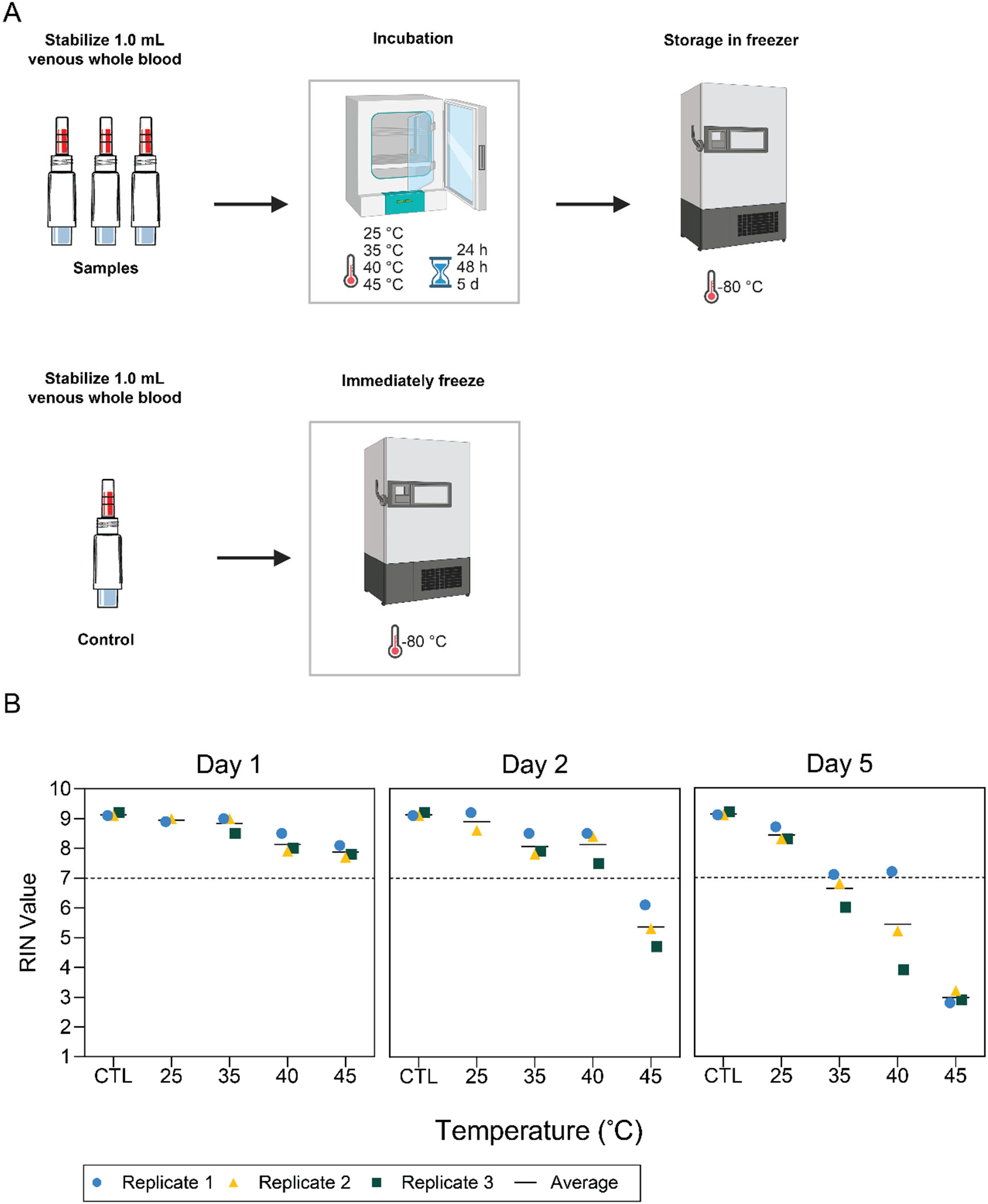
High RIN values are observed at temperatures representing standard shipping conditions. A) Schematic of experimental conditions. homeRNA-stabilized blood samples were incubated at either 25 °C, 35 °C, 40 °C, or 45 °C for 24 hours, 48 hours, and 5 days. Additional samples were also immediately frozen at -80 °C as a control. Created using BioRender.com. B) RNA integrity of extracted samples. Each condition was conducted in triplicate; different symbols represent replicates and bars represent the average of the 3 replicates. One of the replicates for 25 °C data for Day 1 and Day 2 is reported in duplicate due to an error in the RNA extraction process.

### Blood samples stabilized using homeRNA can be stored at temperatures up to 20 °C for up to four weeks with minimal impact on RNA integrity

Different laboratories might have different needs and capabilities in terms of sample storage conditions. It is therefore valuable to evaluate the homeRNA platform’s ability to maintain sample quality when stored at different temperatures (Fig. 8A). We chose any temperature that might be used for sample storage, including room temperature, refrigeration at 4 °C, freezing at - 20 °C, and freezing at -80 °C. We also chose to store stabilized samples for 1 week to represent short-term storage, and 4 weeks to represent a longer-term storage option. We also tested Axygen tubes with its associated cap (i.e., without the homeRNA adapter) and microcentrifuge tubes that are certified for -80 °C storage to compare the homeRNA platform and the Axygen tube’s performances to a gold standard.

**Figure 8.**
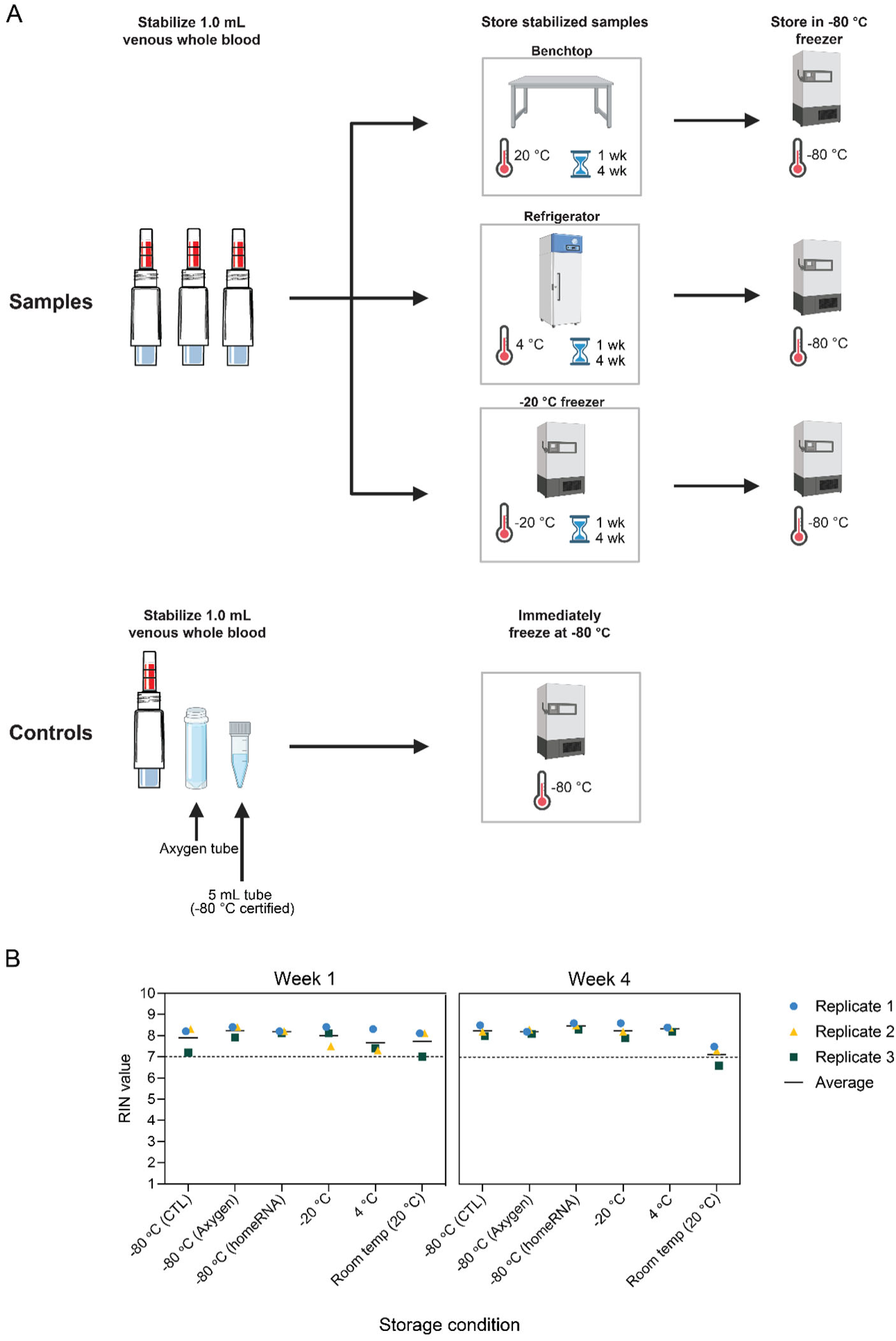
High RIN values are observed across a range of sample cold and ambient storage conditions after four weeks. A) Schematic of experiment. homeRNA-stabilized blood samples were either placed on a benchtop at ambient temperature (20 °C) or stored in a 4 °C refrigerator, -20 °C freezer, or -80 °C freezer. An Axygen tube with its corresponding cap (i.e., without the homeRNA adapter) and a 5 mL microcentrifuge tube certified for -80 °C storage were also used for blood stabilization and immediately stored at -80 °C as additional controls for comparison. Made using BioRender.com. B) RNA integrity of extracted samples. Different symbols represent replicates and bars represent the average of the three replicates. Each condition was performed in triplicate.

All samples stored at 4 °C, -20 °C, and -80 °C for either one week or four weeks had RIN values greater than 7. Additionally, homeRNA samples stored at ambient room temperature (approximately 20 °C) for one week had an average RIN greater than 7 (Fig. 8B). The four-week storage at room temperature is at or near a RIN of 7 and could be used for downstream transcriptomic analysis. It is worth noting that laboratories might want to store samples for longer than four weeks; future work could add additional time points beyond four weeks to continue to test long-term storage conditions. For RNA integrity and yield data for this experiment, see Table S5.

## Conclusion

Recent design modifications to the homeRNA platform make improvements upon the prior versions to increase accessibility and usability for laboratories and to allow for higher-throughput manufacturing. The new platform can be stored for at least three months prior to use, is unaffected by variable sample blood volumes from participants, is robust to climate conditions and additional shaking for durations representing standard shipping conditions, and stabilized blood samples can be stored for at least 4 weeks in any standard laboratory storage condition while still preserving high quality RNA. Future work will continue to test the limits of the homeRNA platform’s ability to withstand extreme conditions for long durations of time, including longer storage times before sampling, longer durations of extreme climates, and longer durations of stabilized sample storage.

## Acknowledgments

This project has been funded in whole or in part with Federal funds from the National Institute on Aging, National Institutes of Health, Department of Health and Human Services, under Contract No. 75N95024C00031. This publication was also supported by the National Center for Advancing Translational Sciences TL1TR002318 (ME, trainee). The REDCap used for human subjects consent and enrollment is supported by the Institute of Translational Health Sciences, which is funded by the National Center for Advancing Translational Sciences of the National Institutes of Health under award number UL1TR002319. The content is solely the responsibility of the authors and does not necessarily represent the official views of the National Institutes of Health. Figures partially created in BioRender. Eakman, M. (2026) https://BioRender.com/l5d7bfi. We would also like to thank the participants for their valued contributions.

## Author contributions

ME, IHR, EB, ST, and ABT conceptualized the validation experiments. DJG designed the modified homeRNA device in CAD and tested prototypes of the updated design. ME and IHR performed first iterations of the validation experiments, including data analysis and associated visuals and write-ups. ME, XW, FS, and TC completed final validation experiments reported in this manuscript. KNA wrote the IRB protocol for venous blood draws performed in this work, and KNA and IHR advised on human subjects study implementation. ME and LGB wrote sections of the manuscript. ME created all visual work presented in the paper. LGB, ST, and ABT advised on study design and execution. All authors revised and approved the manuscript.

## References

(1) Ivanov, S. M.; Tarasova, O. A.; Poroikov, V. V. Transcriptome-Based Analysis of Human Peripheral Blood Reveals Regulators of Immune Response in Different Viral Infections. Front. Immunol. 2023, 14, 1199482. 10.3389/fimmu.2023.1199482

(2) Chaussabel, D. Assessment of Immune Status Using Blood Transcriptomics and Potential Implications for Global Health. Semin. Immunol. 2015, 27 (1), 58–66. 10.1016/j.smim.2015.03.002

(3) Mohr, S.; Liew, C.-C. The Peripheral-Blood Transcriptome: New Insights into Disease and Risk Assessment. Trends Mol. Med. 2007, 13 (10), 422–432. 10.1016/j.molmed.2007.08.003

(4) Yurkovich, J. T.; Hood, L. Blood Is a Window into Health and Disease. Clin. Chem. 2019, 65 (10), 1204–1206. 10.1373/clinchem.2018.299065

(5) Krasowski, M. D. Remote Blood Collection Devices Improve Study Participation from Hard to Reach Populations. J. Appl. Lab. Med. 2024, 9 (5), 874–876. 10.1093/jalm/jfae075

(6) Bybjerg-Grauholm, J.; Hagen, C. M.; Khoo, S. K.; Johannesen, M. L.; Hansen, C. S.; Bækvad-Hansen, M.; Christiansen, M.; Hougaard, D. M.; Hollegaard, M. V. RNA Sequencing of Archived Neonatal Dried Blood Spots. Mol. Genet. Metab. Rep. 2017, 10, 33–37. 10.1016/j.ymgmr.2016.12.004

(7) McDade, T. W.; M. Ross, K.; L. Fried, R.; Arevalo, J. M. G.; Ma, J.; Miller, G. E.; Cole, S. W. Genome-Wide Profiling of RNA from Dried Blood Spots: Convergence with Bioinformatic Results Derived from Whole Venous Blood and Peripheral Blood Mononuclear Cells. Biodemography Soc. Biol. 2016, 62 (2), 182–197. 10.1080/19485565.2016.1185600

(8) Heneghan, N.; Fu, J.; Pritchard, J.; Payton, M.; Allen, R. W. The Effect of Environmental Conditions on the Rate of RNA Degradation in Dried Blood Stains. Forensic Sci. Int. Genet. 2021, 51, 102456. 10.1016/j.fsigen.2020.102456

(9) Haack, A. J.; Lim, F. Y.; Kennedy, D. S.; Day, J. H.; Adams, K. N.; Lee, J. J.; Berthier, E.; Theberge, A. B. *Home* RNA: A Self-Sampling Kit for the Collection of Peripheral Blood and Stabilization of RNA. Anal. Chem. 2021, 93 (39), 13196–13203. 10.1021/acs.analchem.1c02008

(10) Eakman, M.; Stefanovic, F.; Berthier, J.; Robertson, I. H.; Knudsen, L. A.; Cardoso Carvalho, C.; Chen, K.; Atkeson, J.; Eichman, S.; Nguyen, S. H.; Craig, C. A.; Leong, K. M.; Chan, D. W. H.; Adams, K. N.; Olanrewaju, A. O.; Thongpang, S.; Nicholson, T. M.; Haberman, R. H.; Scher, J. U.; Berthier, E.; Theberge, A. B.; Haack, A. J. To the homeRNAmax: Developing an Improved Blood Self-Collection and Stabilization Platform for Remote Transcriptomic Studies. Anal. Chem. 2026, 98 (25), 18363–18372. 10.1021/acs.analchem.5c07772

(11) Eakman, M.; Stefanovic, F.; Liang, W.; Robertson, I. H.; Jindal, A.; Adams, K. N.; Berthier, E.; Haack, A. J.; Theberge, A. B. homeRNA Just Dropped! Streamlining Remote Blood Collection and RNA Stabilization for Smaller Sample Volumes with homeRNAdrop. Bioengineering September 23, 2026. 10.64898/2026.09.22.753625

(12) Brown, L. G.; Haack, A. J.; Kennedy, D. S.; Adams, K. N.; Stolarczuk, J. E.; Takezawa, M. G.; Berthier, E.; Thongpang, S.; Lim, F. Y.; Chaussabel, D.; Garand, M.; Theberge, A. B. At-Home Blood Collection and Stabilization in High Temperature Climates Using homeRNA. Front. Digit. Health 2022, 4, 903153. 10.3389/fdgth.2022.903153

(13) Stefanovic, F.; Brown, L. G.; MacDonald, J.; Bammler, T.; Rinchai, D.; Nguyen, S.; Zeng, Y.; Shinkawa, V.; Adams, K.; Chaussabel, D.; Berthier, E.; Haack, A. J.; Theberge, A. B. Your Blood Is Out for Delivery: Considerations of Shipping Time and Temperature on Degradation of RNA from Stabilized Whole Blood. Anal. Chem. 2025, 97 (3), 1635–1644. 10.1021/acs.analchem.4c04591

(14) Haack, A. J.; Brown, L. G.; Zeng, Y.; Khan, T.; Robertson, I. H.; Kennedy, D. S.; Adams, K. N.; MacDonald, J. W.; Bammler, T. K.; Stefanovic, F.; Moloney, K.; Stolarczuk, J. E.; Alizai, M. Y.; Hassan, G. W.; Lim, F. Y.; Chaussabel, D.; Walker, E. G.; Berthier, E.; Theberge, A. B. A Flexible and Responsive Remote Study Design to Assess Gene Expression Changes During Wildfire Smoke Exposure with homeRNA, an At-Home Blood Sampling Kit.

(15) Stefanovic, F.; Robertson, I.; Moloney, K.; Edelson, J.; Nguyen, S.; Shinkawa, V.; Uchimura, K.; Lin, A.; Le, L.; Tokihiro, J. C.; Takezawa, M. G.; Phan, D.; Schiffer, J. T.; Boeckh, M.; Adams, N.; Waghmare, A.; Errett, N. A.; Berthier, E.; Lim, F. Y.; Theberge, A. B. Research In Your Mailbox: Remote Blood Self-Sampling Enables Participation of Underserved Populations in Longitudinal Studies.

(16) Lim, F. Y.; Kim, S.-Y.; Kulkarni, K. N.; Blazevic, R. L.; Kimball, L. E.; Lea, H. G.; Haack, A. J.; Gower, M. S.; Stevens-Ayers, T.; Starita, L. M.; Boeckh, M.; Hyrien, O.; Schiffer, J. T.; Theberge, A. B.; Waghmare, A. High-Frequency Home Self-Collection of Capillary Blood Correlates IFI27 Expression Kinetics with SARS-CoV-2 Viral Clearance. J. Clin. Invest. 2023, 133 (23), e173715. 10.1172/JCI173715

(17) Lim, F. Y.; Lea, H. G.; Dostie, A. M.; Kim, S.-Y.; van Neel, T. L.; Hassan, G. W.; Takezawa, M. G.; Starita, L. M.; Adams, K. N.; Boeckh, M.; Schiffer, J. T.; Hyrien, O.; Waghmare, A.; Berthier, E.; Theberge, A. B. homeRNA Self-Blood Collection Enables High-Frequency Temporal Profiling of Presymptomatic Host Immune Kinetics to Respiratory Viral Infection: A Prospective Cohort Study. EBioMedicine 2025, 112, 105531. 10.1016/j.ebiom.2024.105531

(18) Jiang, Z.; Lu, Y.; Shi, M.; Li, H.; Duan, J.; Huang, J. Effects of Storage Temperature, Storage Time, and Hemolysis on the RNA Quality of Blood Specimens: A Systematic Quantitative Assessment. Heliyon 2023, 9 (6), e16234. 10.1016/j.heliyon.2023.e16234

(19) Duale, N.; Lipkin, W. I.; Briese, T.; Aarem, J.; Rønningen, K. S.; Aas, K. K.; Magnus, P.; Harbak, K.; Susser, E.; Brunborg, G. Long-Term Storage of Blood RNA Collected in RNA Stabilizing Tempus Tubes in a Large Biobank – Evaluation of RNA Quality and Stability. BMC Res. Notes 2014, 7 (1), 633. 10.1186/1756-0500-7-633

(20) Sarathkumara, Y. D.; Browne, D. J.; Kelly, A. M.; Pattinson, D. J.; Rush, C. M.; Warner, J.; Proietti, C.; Doolan, D. L. The Effect of Tropical Temperatures on the Quality of RNA Extracted from Stabilized Whole-Blood Samples.

(21) Weber, D. G.; Casjens, S.; Rozynek, P.; Lehnert, M.; Zilch-Schöneweis, S.; Bryk, O.; Taeger, D.; Gomolka, M.; Kreuzer, M.; Otten, H.; Pesch, B.; Johnen, G.; Brüning, T. Assessment of mRNA and microRNA Stabilization in Peripheral Human Blood for Multicenter Studies and Biobanks. Biomark. Insights 2010, 5, BMI.S5522. 10.4137/BMI.S5522

(22) Huang, L.-H.; Lin, P.-H.; Tsai, K.-W.; Wang, L.-J.; Huang, Y.-H.; Kuo, H.-C.; Li, S.-C. The Effects of Storage Temperature and Duration of Blood Samples on DNA and RNA Qualities. PLOS ONE 2017, 12 (9), e0184692. 10.1371/journal.pone.0184692

(23) Mueller, O.; Lightfoot, S.; Schroeder, A. RNA Integrity Number (RIN) - Standardization of RNA Quality Control, 2016. https://www.agilent.com/cs/library/applications/5989-1165EN.pdf?srsltid=AYCX9vKpbOfW6xu5oX3X54yRYDjsUy6nMJt15cCNv1kDECVTDQTD8F-t

(24) Schroeder, A.; Mueller, O.; Stocker, S.; Salowsky, R.; Leiber, M.; Gassmann, M.; Lightfoot, S.; Menzel, W.; Granzow, M.; Ragg, T. The RIN: An RNA Integrity Number for Assigning Integrity Values to RNA Measurements. BMC Mol. Biol. 2006, 7 (1), 3. 10.1186/1471-2199-7-3

(25) Gallego Romero, I.; Pai, A. A.; Tung, J.; Gilad, Y. RNA-Seq: Impact of RNA Degradation on Transcript Quantification. BMC Biol. 2014, 12 (1), 42. 10.1186/1741-7007-12-42

(26) Ozsolak, F.; Milos, P. M. RNA Sequencing: Advances, Challenges and Opportunities. Nat. Rev. Genet. 2011, 12 (2), 87–98. 10.1038/nrg2934

(27) Schuierer, S.; Carbone, W.; Knehr, J.; Petitjean, V.; Fernandez, A.; Sultan, M.; Roma, G. A Comprehensive Assessment of RNA-Seq Protocols for Degraded and Low-Quantity Samples. BMC Genomics 2017, 18 (1), 442. 10.1186/s12864-017-3827-y

(28) Jang, J. S.; Holicky, E.; Lau, J.; McDonough, S.; Mutawe, M.; Koster, M. J.; Warrington, K. J.; Cuninngham, J. M. Application of the 3′ mRNA-Seq Using Unique Molecular Identifiers in Highly Degraded RNA Derived from Formalin-Fixed, Paraffin-Embedded Tissue. BMC Genomics 2021, 22 (1), 759. 10.1186/s12864-021-08068-1

