## Supplementary Information for "From Fill to Freeze: Redesigning homeRNA to Enable Robust Sampling and Scalable Downstream Processing Workflows for Remote Blood Transcriptomics"

**Table of Contents:**


| | |
|---|---|
| **Figure S1:** homeRNA tube RNAlater evaporation at room temperature | S1 |
| **Figure S2:** homeRNA tube RNA later evaporation at elevated temperature | S1 |
| **Table S1:** Summary data table for homeRNA aging experiment (shelf life) | S2-S3 |
| **Table S2:** Summary data table for blood:RNAlater ratio experiment | S4 |
| **Table S3:** Summary data table for humidity and shaking experiments | S5 |
| **Table S4:** Summary data table for high temperature stability experiment | S6-S7 |
| **Table S5:** Summary data table for sample storage condition experiment | S8-S9 |

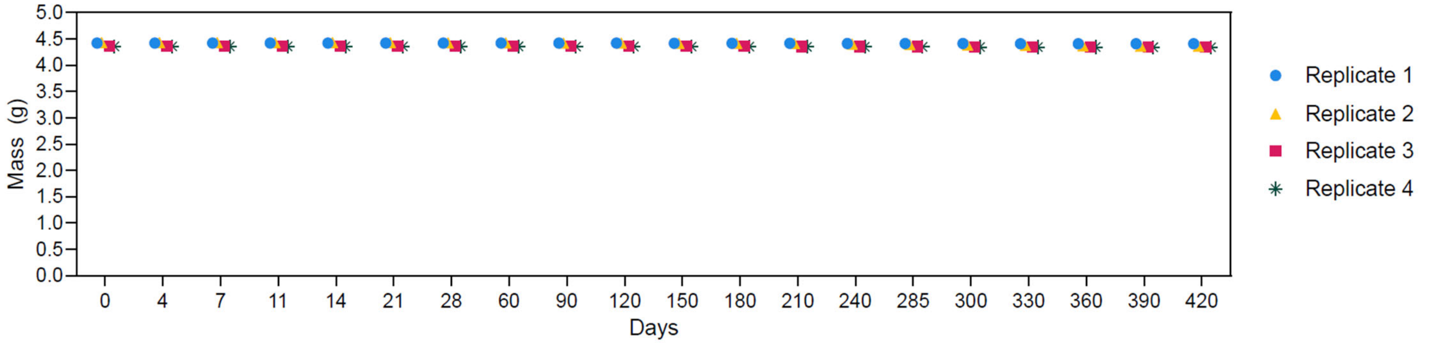


**Figure S1.** homeRNA tube RNAlater evaporation at room temperature. homeRNA tubes were filled with 3.6 mL RNAlater and stored at ambient room temperature (~15.5 ºC - 23.0 ºC) and were weighed after four, seven, eleven, fourteen, and twenty-eight days, then were weighed ~monthly thereafter through day 420. There was minimal evaporation from day zero to day 420 at room temperature.

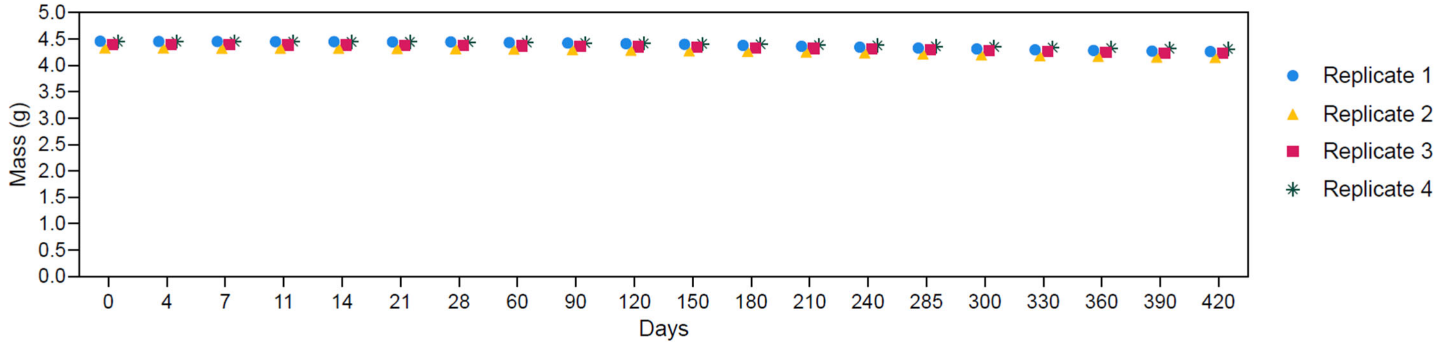


**Figure S2.** homeRNA tube RNAlater evaporation at elevated temperature. homeRNA tubes were filled with 3.6 mL RNAlater and stored in an incubator set to 35 ºC and were weighed after four, seven, eleven, fourteen, and twenty-eight days, then were weighed ~monthly thereafter through day 420. There was approximately 3.850% loss from day zero to day 420. We note that the incubator used for this experiment was shut off in error for up to two weeks between the day 60 and day 90 timepoints.

**Table S1.** Summary data table for the conditions, yield, and RNA Integrity Number (RIN) for the experiment assessing the impact of homeRNA tube aging on RNA integrity (Shelf life - Fig. 3). Blood volume for all conditions is 1 mL.

| Tube storage temperature | Sample incubation temperature | Storage time | Replicate | Yield (µg) | RIN |
|---|---|---|---|---|---|
| Control | -80 °C | Fresh (1 month) | 1 | 3.89 | 7.4 |
| Control | -80 °C | Fresh (1 month) | 2 | 4.97 | 7.6 |
| Control | -80 °C | Fresh (1 month) | 3 | 4.73 | 7.7 |
| 25 ºC | -80 °C | 1 month | 1 | 5.13 | 7.3 |
| 25 ºC | -80 °C | 1 month | 2 | 6.52 | 7.3 |
| 25 ºC | -80 °C | 1 month | 3 | 4.98 | 8.0 |
| 40 ºC | -80 °C | 1 month | 1 | 4.43 | 7.1 |
| 40 ºC | -80 °C | 1 month | 2 | 5.32 | 7.4 |
| 40 ºC | -80 °C | 1 month | 3 | 3.73 | 7.5 |
| Control | 25 ºC | Fresh (1 month) | 1 | 5.85 | 6.6 |
| Control | 25 ºC | Fresh (1 month) | 2 | 4.30 | 7.5 |
| Control | 25 ºC | Fresh (1 month) | 3 | 5.16 | 7.7 |
| 25 ºC | 25 ºC | 1 month | 1 | 6.41 | 7.1 |
| 25 ºC | 25 ºC | 1 month | 2 | 3.54 | 7.7 |
| 25 ºC | 25 ºC | 1 month | 3 | 6.23 | 7.5 |
| 40 ºC | 25 ºC | 1 month | 1 | 2.82 | 5.9 |
| 40 ºC | 25 ºC | 1 month | 2 | 3.81 | 7.3 |
| 40 ºC | 25 ºC | 1 month | 3 | 4.45 | 7.7 |
| Control | 40 ºC | Fresh (1 month) | 1 | 4.04 | 5.9 |
| Control | 40 ºC | Fresh (1 month) | 2 | 4.54 | 6.4 |
| Control | 40 ºC | Fresh (1 month) | 3 | 6.25 | 5.5 |
| 25 ºC | 40 ºC | 1 month | 1 | 3.90 | 6.6 |
| 25 ºC | 40 ºC | 1 month | 2 | 4.27 | 6.4 |
| 25 ºC | 40 ºC | 1 month | 3 | 4.79 | 6.7 |
| 40 ºC | 40 ºC | 1 month | 1 | 3.70 | 7.2 |
| 40 ºC | 40 ºC | 1 month | 2 | 4.25 | 6.3 |
| 40 ºC | 40 ºC | 1 month | 3 | 3.48 | 6.5 |
| Control | -80 °C | Fresh (3 month) | 1 | 4.13 | 8.2 |
| Control | -80 °C | Fresh (3 month) | 2 | 4.18 | 8.2 |

| | | | | | |
|---|---|---|---|---|---|
| Control | -80 °C | Fresh (3 month) | 3 | 4.45 | 8.0 |
| 25 ºC | -80 °C | 3 month | 1 | 5.46 | 8.2 |
| 25 ºC | -80 °C | 3 month | 2 | 4.63 | 8.0 |
| 25 ºC | -80 °C | 3 month | 3 | 4.84 | 7.9 |
| 40 ºC | -80 °C | 3 month | 1 | 4.46 | 7.2 |
| 40 ºC | -80 °C | 3 month | 2 | 4.72 | 7.2 |
| 40 ºC | -80 °C | 3 month | 3 | 5.28 | 7.1 |
| Control | 25 ºC | Fresh (3 month) | 1 | 5.63 | 7.9 |
| Control | 25 ºC | Fresh (3 month) | 2 | 5.73 | 8.3 |
| Control | 25 ºC | Fresh (3 month) | 3 | 5.59 | 7.9 |
| 25 ºC | 25 ºC | 3 month | 1 | 5.04 | 7.9 |
| 25 ºC | 25 ºC | 3 month | 2 | 4.71 | 8.1 |
| 25 ºC | 25 ºC | 3 month | 3 | 4.74 | 7.8 |
| 40 ºC | 25 ºC | 3 month | 1 | 4.12 | 8.2 |
| 40 ºC | 25 ºC | 3 month | 2 | 4.04 | 8.1 |
| 40 ºC | 25 ºC | 3 month | 3 | 4.23 | 7.8 |
| Control | 40 ºC | Fresh (3 month) | 1 | 4.02 | 6.7 |
| Control | 40 ºC | Fresh (3 month) | 2 | 4.57 | 7.2 |
| Control | 40 ºC | Fresh (3 month) | 3 | 4.32 | 6.5 |
| 25 ºC | 40 ºC | 3 month | 1 | 3.89 | 6.3 |
| 25 ºC | 40 ºC | 3 month | 2 | 3.93 | 6.4 |
| 25 ºC | 40 ºC | 3 month | 3 | 4.93 | 6.6 |
| 40 ºC | 40 ºC | 3 month | 1 | 4.74 | 6.3 |
| 40 ºC | 40 ºC | 3 month | 2 | 4.11 | 5.9 |
| 40 ºC | 40 ºC | 3 month | 3 | 4.12 | 5.8 |

**Table S2.** Summary data table for the conditions, yield, and RNA Integrity Number (RIN) for the experiment assessing whether excess RNAlater (i.e., blood:RNAlater ratio - Fig. 4) impacts RNA integrity.

| Blood Volume | Temperature | Replicate | Yield (µg) | RIN |
|---|---|---|---|---|
| 0.25 mL | 25 ºC | 1 | 2.28 | 7.9 |
| 0.25 mL | 25 ºC | 2 | 2.12 | 7.8 |
| 0.50 mL | 25 ºC | 1 | 3.78 | 7.6 |
| 0.50 mL | 25 ºC | 2 | 2.40 | 7.8 |
| 1.00 mL | 25 ºC | 1 | 6.52 | 7.9 |
| 1.00 mL | 25 ºC | 2 | 4.79 | 7.7 |
| 0.25 mL | -80 ºC | 1 | 1.99 | 8.0 |
| 0.25 mL | -80 ºC | 2 | 0.92 | 8.2 |
| 0.50 mL | -80 ºC | 1 | 3.14 | 7.6 |
| 0.50 mL | -80 ºC | 2 | 2.57 | 8.0 |
| 1.00 mL | -80 ºC | 1 | 5.76 | 7.8 |
| 1.00 mL | -80 ºC | 2 | 4.66 | 7.8 |

**Table S3.** Summary data table for the conditions, yield, and RNA Integrity Number (RIN) for the experiments assessing the impact of humidity and shaking on RNA integrity (Fig. 5-6). Blood volume for all conditions is 1 mL.

| Condition | Temperature | Replicate | Yield (μg) | RIN |
|---|---|---|---|---|
| Ambient Humidity (~35%) | 25 ºC | 1 | 4.8 | 8.5 |
| Ambient Humidity (~35%) | 25 ºC | 2 | 4.5 | 8.9 |
| Ambient Humidity (~35%) | 25 ºC | 3 | 5.2 | 8.7 |
| Low Humidity (~10%) | 25 ºC | 1 | 6.0 | 8.6 |
| Low Humidity (~10%) | 25 ºC | 2 | 5.1 | 8.3 |
| Low Humidity (~10%) | 25 ºC | 3 | 5.7 | 8.8 |
| High Humidity (~90%) | 25 ºC | 1 | 4.5 | 8.7 |
| High Humidity (~90%) | 25 ºC | 2 | 5.0 | 8.2 |
| High Humidity (~90%) | 25 ºC | 3 | 5.6 | 8.8 |
| Vibration | 25 ºC | 1 | 6.0 | 9.0 |
| Vibration | 25 ºC | 2 | 5.9 | 8.8 |
| Vibration | 25 ºC | 3 | 5.8 | 8.5 |
| No Vibration | 25 ºC | 1 | 6.1 | 8.9 |
| No Vibration | 25 ºC | 2 | 5.3 | 8.8 |
| No Vibration | 25 ºC | 3 | 6.0 | 8.7 |
| Control | -80 ºC | 1 | 6.2 | 9.0 |
| Control | -80 ºC | 2 | 5.6 | 8.6 |
| Control | -80 ºC | 3 | 5.3 | 8.9 |

**Table S4.** Summary data table for the conditions, yield, and RNA Integrity Number (RIN) for the experiment assessing the impact of high temperatures on RNA integrity (Fig. 7). Blood volume for all conditions is 1 mL.

| Temperature | Incubation Time | Replicate | Yield (μg) | RIN |
|---|---|---|---|---|
| 25 ºC | 24 h | 1 | 5.37 | 8.9 |
| 25 ºC | 24 h | 2 | 4.04 | 9.0 |
| 25 ºC | 24 h | 3 | N/A | N/A |
| 25 ºC | 48 h | 1 | 5.11 | 9.2 |
| 25 ºC | 48 h | 2 | 3.47 | 8.6 |
| 25 ºC | 48 h | 3 | N/A | N/A |
| 25 ºC | 5 d | 1 | 5.68 | 8.7 |
| 25 ºC | 5 d | 2 | 4.18 | 8.3 |
| 25 ºC | 5 d | 3 | 5.03 | 8.3 |
| 35ºC | 24 h | 1 | 5.47 | 9.0 |
| 35ºC | 24 h | 2 | 3.97 | 9.0 |
| 35ºC | 24 h | 3 | 4.60 | 8.5 |
| 35ºC | 48 h | 1 | 5.03 | 8.5 |
| 35ºC | 48 h | 2 | 3.60 | 7.8 |
| 35ºC | 48 h | 3 | 3.81 | 7.9 |
| 35ºC | 5 d | 1 | 5.39 | 7.1 |
| 35ºC | 5 d | 2 | 4.44 | 6.8 |
| 35ºC | 5 d | 3 | 4.74 | 6.0 |
| 40 ºC | 24 h | 1 | 6.46 | 8.5 |
| 40 ºC | 24 h | 2 | 3.43 | 7.9 |
| 40 ºC | 24 h | 3 | 4.17 | 8.0 |
| 40 ºC | 48 h | 1 | 5.18 | 8.5 |
| 40 ºC | 48 h | 2 | 4.26 | 8.4 |
| 40 ºC | 48 h | 3 | 3.83 | 7.5 |
| 40 ºC | 5 d | 1 | 4.95 | 7.2 |
| 40 ºC | 5 d | 2 | 3.04 | 5.2 |
| 40 ºC | 5 d | 3 | 4.32 | 3.9 |
| 45 ºC | 24 h | 1 | 5.56 | 8.1 |
| 45 ºC | 24 h | 2 | 3.44 | 7.7 |
| 45 ºC | 24 h | 3 | 4.91 | 7.8 |

| | | | | |
|---|---|---|---|---|
| 45 ºC | 48 h | 1 | 5.04 | 6.1 |
| 45 ºC | 48 h | 2 | 3.31 | 5.3 |
| 45 ºC | 48 h | 3 | 4.69 | 4.7 |
| 45 ºC | 5 d | 1 | 4.66 | 2.8 |
| 45 ºC | 5 d | 2 | 4.13 | 3.2 |
| 45 ºC | 5 d | 3 | 4.50 | 2.9 |
| -80 ºC | | 1 | 5.24 | 9.1 |
| -80 ºC | | 2 | 4.57 | 9.1 |
| -80 ºC | | 3 | 4.72 | 9.2 |

**Table S5.** Summary data table for the conditions, yield, and RNA Integrity Number (RIN) for the experiment assessing the impact of sample storage conditions on RNA integrity (Fig. 8). Blood volume for all conditions is 1 mL.

| Temperature | Tube used | Incubation Time | Replicate | Yield (μg) | RIN |
|---|---|---|---|---|---|
| -80 °C | Control | 1 week | 1 | 4.84 | 8.2 |
| -80 °C | Control | 1 week | 2 | 4.99 | 8.3 |
| -80 °C | Control | 1 week | 3 | 3.94 | 7.2 |
| -80 °C | Control | 4 week | 1 | 3.57 | 8.5 |
| -80 °C | Control | 4 week | 2 | 3.84 | 8.2 |
| -80 °C | Control | 4 week | 3 | 3.92 | 8.0 |
| -80 °C | Axygen tube | 1 week | 1 | 4.32 | 8.4 |
| -80 °C | Axygen tube | 1 week | 2 | 4.59 | 8.4 |
| -80 °C | Axygen tube | 1 week | 3 | 3.87 | 7.9 |
| -80 °C | Axygen tube | 4 week | 1 | 3.68 | 8.2 |
| -80 °C | Axygen tube | 4 week | 2 | 4.09 | 8.3 |
| -80 °C | Axygen tube | 4 week | 3 | 4.33 | 8.1 |
| -80 °C | homeRNA tube | 1 week | 1 | 4.53 | 8.2 |
| -80 °C | homeRNA tube | 1 week | 2 | 4.93 | 8.2 |
| -80 °C | homeRNA tube | 1 week | 3 | 3.12 | 8.1 |
| -80 °C | homeRNA tube | 4 week | 1 | 4.15 | 8.6 |
| -80 °C | homeRNA tube | 4 week | 2 | 4.24 | 8.5 |
| -80 °C | homeRNA tube | 4 week | 3 | 4.15 | 8.3 |
| Room temp (~20 °C) | homeRNA tube | 1 week | 1 | 4.34 | 8.1 |
| Room temp (~20 °C) | homeRNA tube | 1 week | 2 | 4.42 | 8.1 |
| Room temp (~20 °C) | homeRNA tube | 1 week | 3 | 3.44 | 7.0 |
| Room temp (~20 °C) | homeRNA tube | 4 week | 1 | 6.68 | 7.5 |
| Room temp (~20 °C) | homeRNA tube | 4 week | 2 | 5.30 | 7.3 |
| Room temp (~20 °C) | homeRNA tube | 4 week | 3 | 5.57 | 6.6 |
| 4 °C | homeRNA tube | 1 week | 1 | 3.65 | 8.3 |
| 4 °C | homeRNA tube | 1 week | 2 | 4.90 | 7.3 |
| 4 °C | homeRNA tube | 1 week | 3 | 2.92 | 7.4 |
| 4 °C | homeRNA tube | 4 week | 1 | 4.26 | 8.4 |
| 4 °C | homeRNA tube | 4 week | 2 | 3.30 | 8.4 |

| | | | | | |
|---|---|---|---|---|---|
| 4 °C | homeRNA tube | 4 week | 3 | 4.38 | 8.2 |
| -20 °C | homeRNA tube | 1 week | 1 | 4.05 | 8.4 |
| -20 °C | homeRNA tube | 1 week | 2 | 4.91 | 7.5 |
| -20 °C | homeRNA tube | 1 week | 3 | 2.85 | 8.1 |
| -20 °C | homeRNA tube | 4 week | 1 | 4.02 | 8.6 |
| -20 °C | homeRNA tube | 4 week | 2 | 4.58 | 8.2 |
| -20 °C | homeRNA tube | 4 week | 3 | 4.19 | 7.9 |
